# Socioecological differentiation and the evolution of brain size and synaptic architecture in predatory ants, *Neoponera*

**DOI:** 10.64898/2026.06.27.735026

**Authors:** Frank Azorsa, James F.A. Traniello

## Abstract

Brain size and structure are hypothesized to be adaptively designed to satisfy the behavioral requirements of securing food and living socially. The importance of these socioecological and sociobiological selective forces in brain evolution is constantly debated. Socioecological divergence is striking in the Neotropical ant genus *Neoponera*: *N*. *apicalis* is a generalist solitary predator forming small colonies of ∼100 whereas *N*. *commutata* colonies are approximately 10 times as large and workers pheromonally organize cooperatively raids only on *Syntermes* termite colonies. We interspecifically compared the size and structure of the compound eyes, size and number of antennal glomeruli, mosaic brain scaling and synaptic processing (microglomeruli-MG). Our results indicate that *N*. *apicalis* workers have a larger number of ommatidia, antennal lobe glomeruli, and allometrically larger antennal and optic lobes than *N*. *commutata*. These sensory traits were associated with differences in higher-order processing architectures in the mushroom body (MB) microglomeruli (MG). *N*. *commutata* workers had an allometrically larger MB, perhaps due to their socially complex chemical foraging communication, although MG density in *N*. *apicalis* was higher in both the MB lip and collar, regions associated with processing olfactory and visual information, respectively. The increase in MG density in *N*. *apicalis* may be associated with higher demands for navigation, learning, and memory, as well as a higher density of antennal lobe glomeruli to support prey odor discrimination. In contrast, *N*. *commutata* workers had larger ommatidia and antennal lobe glomeruli. Larger ommatidia correlate with their diurnal/nocturnal habits and a larger MB Our findings indicate that differences in behavioral performance demands associated with socioecological differentiation are reflected in variation in visual and olfactory system structure, brain size, mosaicism, and synaptic organization. Our results support both social and ecological brain hypothesis as drivers of mosaic brain evolution.

## Introduction

The importance of sociobiological and ecological selective forces on brain evolution is actively debated. The social brain hypothesis predicts that brain size increases in relation to group size and associated complexity of social interactions (Dunbar 1998; Dunbar and Shultz 2017). Alternatively, ecological theories of brain evolution posit that diet is more significant than social complexity, due to the cognitive demands associated with finding and processing food (Clutton and Harvey 1980; DeCasien et al. 2017; Farris and Roberts 2005; Goldman-Huertas et al. 2015; Harvey et al. 1980; Powell et al. 2017; Rosati 2017) and that behavioral and/or cognitive abilities and their neurobiological support systems are needed to adapt to changing environments (Bertrand et al. 2022; Michaud et al. 2022). Brain compartment allometric structure, or mosaicism, appears to adaptively reflect selective demands of social life, ecological adaptation, and life-history traits (Barton and Harvey 2000; Boch et al. 2024; DeCasien and Higham 2019; Finerelli and Flynn 2009; Iwanuk et al. 2004; Maguire et al. 2000; Michaud et al. 2022; Sayol et al. 2020; Shultz and Dunbar 2010, 2022; Sukhum et al. 2018). A role for sociality appears to be supported in some insects. Desert locusts, for example, have larger brains with altered compartmental allometries after they transition from a solitary to gregarious life history stage; however, these neuroanatomical changes correlate not only with social complexity, but also dietary change (Ott and Rogers 2010). Diet also correlates with brain evolution in solitary herbivorous insects (Couto et al. 2023; Farris and Roberts 2005).

The influence of diet and/or sociality on brain evolution is thus not well understood and ongoing controversy concerning the evolutionary pressures that lead to encephalization has extended to eusocial insects. Workers are sterile and selection targets behaviors that enhance colony-level fitness rather than individual reproductive success. This striking difference in the fitness consequences of social behavior between these clades raises questions concerning the application of vertebrate-oriented theories such as the social brain hypothesis to eusocial insects (Chittka and Niven 2009; Farris 2016; Lihoreau et al. 2012, 2019; Traniello et al. 2022). Nevertheless, the basic concepts of social and ecological brain theory provide frameworks to examine the roles of social complexity, diet and foraging ecology, and associated worker behavioral performance demands on brain size and structure. Workers in species characterized by greater complexity in social organization may have larger brain size relative to body size compared to socially basic species (Jaffe and Perez 1989; Kamhi et al. 2016; Wehner et al. 2007). However, brain size may decrease with increased social complexity in phylogenetically remote clades (Desilva et al. 2021; Riveros et al. 2012). In bees, brain size is linked to diet but not sociality (Sayol et al. 2020) and in some ants and wasps, brain size and mosaicism correlate with diet, sensory ecology, morphological caste, task specialization, age, and task plasticity (Muratore et al. 2022; Muscedere and Traniello 2012; O’Donnell et al. 2013; Rosner et al. 2017; Smith et al. 2023).

Ants are remarkably diverse in social structure, diet, and patterns of sensory system and brain evolution (Hölldobler and Wilson 1990; Smith et al. 2023; Traniello et al. 2022). Principal sensory inputs to the brain are provided by the compound eyes and antennae, which detect visual information and chemical signals, respectively (Hansson and Stensmyr 2011; Nilsson and Kelber 2007). Ant brains are composed of compartments that serve distinct behavioral functions. The mushroom bodies (MBs), which are responsible for higher-order processing, learning, and memory, have lip and collar regions that receive olfactory and visual information, respectively (Gronenberg 2008). The optic and antennal lobes (OL, AL) are associated with the perception of visual and olfactory stimuli, respectively. The OL in ants has three regions, the lamina, medulla, and lobula (Gronenberg and Holldobler 1999), responsible for detecting contrast, color and small-field motion, and color and wide-field motion detection, shape, and panoramas, respectively (Dyer et al. 2011; Gronenberg 2008; Strausfeld 1989). The AL is composed of glomeruli that receive olfactory input from antennal sensory neurons (Anton and Homberg 1999). The central complex (CX) is associated with navigation and locomotion (Buehlman et al. 2023; Collet et al. 2025; Strausfeld 2012) and the subesophageal zone (SEZ) provides motor control of the mouthparts (Kendroud et al. 2018) These brain compartments and their functional subdivisions, as well as brain size, scale allometrically with the sensory demands of task performance (Amador-Vargas et al. 2015; Gordon et al. 2017, 2019; Kamhi et al. 2016; Muratore et al. 2022; Muscedere and Traniello 2012; Muscedere et al. 2014; O’Donnell et al. 2018; Riveros et al. 2012; Traniello et al. 2022; Valadares et al 2026).

In addition to adaptive variation in neuropil allocation, worker task performance is underpinned by synaptic circuitry in the brain (e.g., Rossler 2019, 2023; Rossler et al 2023; Seid et al. 2005). The MB lip and collar receive olfactory and visual information, respectively, and neuronal connections in each region are organized in synaptic complexes called microglomeruli (MG; Groh and Rossler 2011). These microprocessors change in density with task-related demands for information processing, learning, and memory (Collet et al. 2025; Falibene et al. 2015; Kamhi et al. 2017; Li et al. 2017) independent of worker body size (Groh et al. 2014). In honeybees and leafcutter ants, the formation of olfactory long-term memory is associated with an increase in MG density (Hourcade et al. 2010; Falibene et al. 2015). Visual experience in foragers of the ant *Cataglypis fortis* correlates with a decrease in MG density in the MB collar (Stieb et al. 2010). Diet, age-related task repertorie expansion (Gordon et al. 2018; Seid et al. 2005), and the level of social complexity appear to influence MG density. An increase in MG volume occurs in older honeybee workers (Groh et al. 2012), indicating an association between synaptic changes (an increase in the size of projection neuron boutons, more synaptic vesicles and ribbon synapses, and a larger number of postsynaptic partners) and behavioral experience. In ant species characterized by highly advanced social organization, MG are larger and denser (Kamhi et al. 2016, 2017). Major workers of the weaver ant *Oecophylla smaragdima,* for example, have more dense and larger MG in older workers, which communicate chemically and navigate visually to a greater extent than minor or newly eclosed workers.

The evolutionary success and ecological dominance of ants can be attributed to their social organization and relationship to dietary diversification (Hölldobler and Wilson 1990; Smith et al. 2023). Many predatory ants are socially ancestral (“primitive”) species that form small colonies, and similar to the earliest ant lineage and their wasp-like ancestors, are highly predaceous (Farris and Schulmeister 2011; Hölldobler and Wilson 1990; Lepeco et al. 2025; Perrichot et al. 2016; Schmidt and Shattuck 2014). In contrast, other species have evolved large colonies and socially complex traits such as worker polymorphism, division of labor, and chemically coordinated foraging (Anderson and McShea 2001; Azorsa et al. 2022; Bell-Roberts et al. 2024). A diverse diet of invertebrates collected by solitary huntresses characterizes the foraging ecology of most predatory ants, but some species specialize on termites, which are also eusocial, form large colonies, and are an energetically valuable clumped food source. The dietary shift from prey randomly distributed in space and time to clumped prey with spatial and temporal persistence involves changes in the behavioral and/or cognitive demands of foraging, resulting in an increase in olfaction during pheromonally organized group predation, whereas solitary huntresses generally rely on vision (Dejean and Lachaud 2011; Downing 1978; Graham and Philippides 2017; Hölldobler and Traniello 1980; Mill 1984).

Ponerine ants are a clade of predatory species characterized by remarkable differentiation in diet and social organization (Azorsa et al. 2022). Clade-restricted comparative analyses of closely related but socioecologically divergent species can enhance our understanding of the relationship of sociality, diet, foraging ecology, worker behavior, brain mosaicism, and synaptic organization (Godfrey and Gronenberg 2019). The difference in colony size, an element of social complexity, and diet is striking in sister species of the Neotropical ant genus *Neoponera*. Workers of *N*. *apicalis* are generalist, diurnal, central-place solitary predators, lack food recruitment communication, and have small colonies (∼100 workers; Schmidt and Shattuck 2014). In contrast, workers of *N. commutata* are dietary specialists, attacking only colonies of the termite *Syntermes* in group raids organized by trail pheromones. *N. commutata* workers attack termite colonies in chemically coordinated groups, have an army-ant like nomadic life cycle, and diurnal and nocturnal activity (Mill 1984). Are pronounced differences in diet, social structure, foraging ecology, and worker behavior associated with adaptive sensory systems, brain mosaicism, and synaptic organization?

We hypothesize that sensory systems, brain size, compartment allometry, relative brain compartment investment, and synaptic organization in *N*. *apicalis* and *N*. *commutata* correlate with colony size, a proxy for social complexity (Anderson and McShea 2001), foraging behavior and diet (Table 1). Specifically, *N*. *apicalis* is predicted to have a higher density of ommatidia and increased downstream investment in the OL, lamina, and MB collar to detect and process visual inputs. If diet has an influence on brain evolution, then workers of *N*. *apicalis* will have a higher density of MG in both the MB-lip and MB-collar because of their greater demands for navigational skills whereas *N*. *commutata* will have larger MG because of the processing of chemical cues essential for communication during foraging and recruitment. In comparison, if colony size increases social interactions that drive brain evolution, then *N*. *commutata* workers should have an allometrically large MB to process social information. If diet and worker foraging behavior influence sensory systems and mosaic structure, then *N*. *commutata* workers are posited to have reduced relative investment in the OLs and MB collar and increased in the size of the AL and MB lip as well as larger AL glomeruli given higher demands for processing olfactory information. We considered that brain size and structure at the tissue and cellular level could result from either diet or sociality.

**Table 1.** Relationships between diet, colony size, (social complexity), foraging ecology and behavior, and brain structure. Hypothesized neural phenotypes, and results are presented. Brain scaling predictions based on the relative importance of diet and cognitive demands for processing social information (reflected in colony size) for dietary generalist and dietary specialist ponerine ants that vary in colony size.

| Species | Diet and colony size | Behavioral requirement | Predicted sensory system and brain structure | Results/statistical significance |
| --- | --- | --- | --- | --- |
| *N. apicalis* | Generalist carnivore/omnivore; small colony size | Greater visual processing for navigation involving learning and memory | Higher density of ommatidia, larger OL, larger MB collar | Higher density of ommatidia, larger OL and MB collar size |
|  |  |  | Allometrically larger CX | Allometrically larger CX |
|  |  |  | Higher density of MG | Higher density of MG in both the lip and the collar |
| *N. commutata* | *Syntermes* specialist carnivore; relatively large colony size | Recognition of olfactory cues specific to *Syntermes* | Reduced optic lobes, MB collar; increased AL size, increased AL glomeruli and MB lip size | Larger mean glomeruli size, and larger MB lip |
|  |  | Greater social interaction during raiding behavior | Allometrically larger MB | Allometrically larger MB |
|  |  | Visual processing for diurnal and low-light activity | Larger ommatidia size, larger lobula and larger MG in the MB collar | Larger ommatidia, lobula, and larger MG in the MB collar |
|  |  | Dependence on chemical signaling during foraging | Larger MG size | Larger MB lip MG |

## Material and methods

### Ant collection and laboratory culture

Workers were sampled from four queenright colonies of each species collected at two field research stations in Peru, located in Loreto and Madre de Dios. The Amazon Conservatory for Tropical Studies (ACTS), and Los Amigos Biological Station, respectively. Laboratory colonies were maintained in an environmental chamber at 25 C°, 65% humidity with a 12h:12h light: dark cycle in test tubes partially filled with water and tightly plugged with cotton placed inside plastic boxes coated with Fluon®. Colonies of *N*. *apicalis* were provisioned *ad lib* every other day with live wingless fruit flies, crickets, mealworms, termites, and sugar water. *N*. *commutata* was provisioned *ad lib* with termites (*Zootermopsis*) each morning. Due to their nomadic life cycle, highly specialized diet, environmental sensitivity and thus difficulty to maintain in culture, worker brains were collected and processed no more than a week after the arrival of colonies in the laboratory.

### Compound eye imaging and structural measurements

Intact compound eyes were imaged to create 3D stacks to measure ommatidia number (ON), and average ommatidial diameter (D) following Arganda et al. (2020). Eyes were stored in 70% ethanol, washed in 100% ethanol (3 × 10 min) before mounting. Extraneous cuticle was removed to allow eyes to lie flat and were then mounted in methyl salicylate and imaged using a Nikon C2Si confocal microscope with a 10X objective. Cuticle has natural fluorescence. Cellpose was used to quantify ommatidia number (ON) (Stringer et al. 2021). Mean ommatidial diameter was calculated from the average diameter of all ommatidia using Image J. The eye surface area was calculated from the mean ommatidial diameter (surface area = ON × π × [0.5 × D]^2^). Ommatidial density (number of ommatidia per surface area unit) was calculated by dividing the number of ommatidia by eye surface area (Arganda et al. 2020).

### Immunohistochemistry, confocal microscopy, and neuroimage analysis

Brains were processed using a modified immunohistochemistry protocol (Gordon et al. 2017). Brains were dissected in ice cold HEPES buffered saline (HBS) and fixed overnight at 4°C on a shaker in 1% zinc formaldehyde. Brains were then washed in HBS (6 x 10 min), transferred to Dent’s fixative (4:1 methanol: dimethyl sulfoxide) for 1 to 2 hours at room temperature, and stored in 100% methanol until further processing. Brains were rehydrated in 0.1M Tris buffer before blocking for 1 hour in a normal goat serum (NGS) solution (PBSTN) (5% NGS + 0.005% sodium azide in 0.2% Triton-X phosphate buffered saline [PBST]). We used a monoclonal *Drosophila* synapsin-1 antibody (anti SYNORF1, AB_528479) purchased from the Developmental Studies Hybridoma Bank (DSHB, University of Iowa, catalog 3C11) as our primary antibody. After blocking, brains were incubated for four nights at 4°C on a shaker in primary antibody, (diluted 1:30 SYNORF1 in PBSTN). Subsequently, brains were washed in 0.2% PBST (6 x 10 min) and incubated for another three nights wrapped in foil at 4°C on a shaker in AlexaFluor488 (ThermoFisher) goat anti-mouse secondary antibody (1:100 in PBSTN). After secondary incubation, brains were washed (6 x 10 min in 0.2% PBST) and dehydrated in a series of ethanol in PBS (10 minutes each in 30, 50, 70, 95, 100, 100%) and stored overnight at -20°C wrapped in foil. Finally, brains were cleared and mounted with methyl salicylate in custom stainless steel well slides. Stained brains were imaged on a Nikon C2Si confocal microscope with a 10X objective and optically sectioned in the horizontal plane (2.8 μm steps). Images were manually segmented in Amira (FEI v 6.0) to generate compartment volumes for one hemisphere of each brain: optic lobe (OL), antennal lobe (AL), mushroom body calyces (MBC), mushroom body peduncle and lobes (MBP), central complex (CX), and subesophageal zone (SEZ), in addition to the rest of the undifferentiated central brain (ROCB). Regions that spanned both hemispheres (CX and SEZ) were traced in full and their volumes were divided by half to equate to one hemisphere. The relative investment in each of six functional subregions defined earlier was calculated by dividing the volume of the region of interest by the volume of the rest of the central brain (ROCB), and the absolute total volume of one hemisphere was calculated as the sum of AL, OL, MBC, MBP, CX, SEZ and ROCB.

### Microglomeruli imaging and quantification

Brains were processed using a modified protocol (Gordon et al. 2018; Kamhi et al. 2016). Brains were dissected in ice-cold HEPES-buffered saline and fixed at 4°C overnight on a rotator in 4% paraformaldehyde in 0.1M PBS. Brains were washed with 0.1 M PBS (3 x 20 min), before embedding in low melting point agarose (5.5g/mL PBS) and cut into 100μm sections using a vibratome. Sections were incubated with 2% and 0.2% PBST each for 20 minutes to permeabilize the tissue before blocking for one hour at room temperature on a plate shaker in 2% normal goat serum (NGS) in 0.2% PBST. Sections were incubated for at least three nights at room temperature while covered in foil on a plate shaker in primary antibody, SYNORF1 (1:50), and AlexaFluor488-phalloidin (1:500). Brain sections were washed in 0.1M PBS (5 x 20 min) and incubated overnight at room temperature while covered in foil on a plate shaker in AlexaFluor568 goat anti- mouse secondary antibody (1:250). Brains sections were incubated overnight in 60% glycerol followed by 80% glycerol for 30 min and then mounted in 80% glycerol on glass slides sealed with nail polish. Prepared slides were imaged (1024 x 1024 pixel resolution) with an oil immersion 60X objective (NA = 1.4) without digital zoom on a Nikon C2Si confocal microscope. Optical sections in a plane which the MB peduncle bisects the MB calyces were made. Each image typically contained both medial and lateral calyces. MG density and size were measured using a modified protocol of Gordon et al. (2018). Two adjacent circles (400μm2 each) were overlaid in the lip region and one circle (400 μm2) on the collar region of each calyx. Cellpose was used to quantify all MG (Stringer et al. 2021) and ImageJ was used to measure the size of all MG in the 400 μm2 area. For each worker, average density of MG in the lip and the collar were calculated by dividing the counts MG by 400 μm2. Similarly, average MG length was calculated for each worker for the lip and the collar region.

### Statistical analysis

Comparisons among workers of *N*. *apicalis* and *N*. *commutata* brain compartment metrics were performed using RStudio. We estimated the intercept and slope for both species by fitting bivariate standardized major axis (SMA) lines to the log-transformed variates using SMATR version 3.4-8 (Warton et al. 2012). We used a log-likelihood test for common slope and Wald tests for common elevation and grade shifts. Each brain compartment (AL, OL, MB, MB peduncle, MB lip, MB collar, CX, and SEZ) was normalized by the ROCB. We first tested for common slopes (β), to test if regression slopes for each species were equal; p-values > 0.05 indicate both species share a common slope. We next tested for allometric scaling, to test if a brain compartment increases in size allometrically or isometrically. We used the scaling exponent (β) to test if a brain compartment is isometric, if β = 1 then the brain compartment increases in size isometrically. We then used the slope index (SI) to compare those exponents between species, an SI = 1 indicates that both species share the same scaling relationship. SI was calculated using the next formula (SI = β*_N. commutata /βN. apicalis_*). Lastly, for comparisons where the two species shared a common scaling slope we calculated a grade shift index (GSI) to test how much larger a brain compartment is in one species relative to another, holding the size of the ROCB constant. If GSI > 1, *N*. *commutata* had a larger brain region, and if GSI < 1, *N*. *apicalis* had a larger brain region. GSI was calculated as 10 to the power of the difference in elevation of both species (10^elevation *N*. *commutata*-elevation *N*. *apicalis*^).

We compared the absolute and relative brain volume (brain volume relative to the head width) and relative investment (brain compartment relative to the ROCB volume) in each functional brain compartment. We normalized each brain compartment by the ROCB volume for scaling analysis. Analysis of covariance (ANCOVA) was used to determine if there were significant differences in absolute and relative brain volume, relative investment in each brain compartment, and the number and size of AL glomeruli and MB microglomeruli.

## Results

### Interspecific differences in absolute and relative brain volume

To quantify neuroarchitectural variation and scaling between species, we compared the absolute and relative brain volume (brain volume as a proportion of head size). Absolute brain volume was larger in *N*. *commutata* workers (p > 0.05) but relative brain volume was not significantly different (*p* > 0.05; Figure 1 d,e). Therefore, absolute brain volume scaled with body size and relative brain volume did not differ interspecifically. We also compared the absolute size of all functional brain compartments: the OL was larger in *N*. *apicalis*, whereas the AL and MB were larger in *N*. *commutata* workers (Figure 1). The absolute size of the CX and the SEZ were not significantly different among both species.

**Figure 1.**
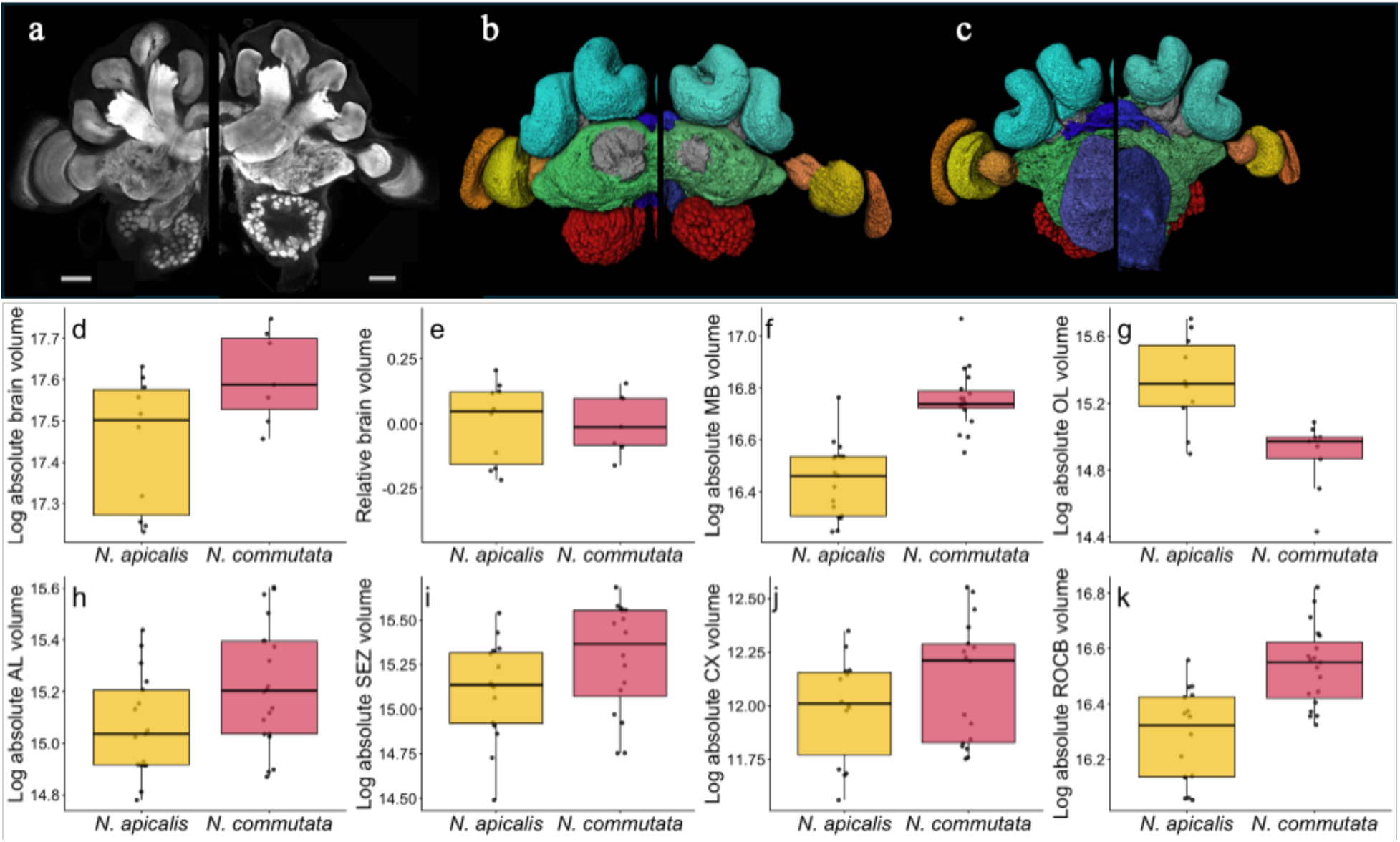
Interspecific comparison of absolute and relative brain size, and absolute size of functional brain compartments in two *Neoponera* ant species. Upper pannel: Confocal images (a) and 3-D reconstructions (b, c) of the brain of *N*. *apicalis* (left) and *N*. *commutata* workers (right). Dorsal view (b), and ventral view (c). Scale bars = 50μm. Lower panel: Absolute (d) and relative brain size (e). Absolute size of invidual brain compartments: MB (f), OL (g), AL (h), SEZ (i), CX (j), and ROCB (k).

### Comparative brain compartment allometry

Scaling relationships between brain compartments were analyzed by fitting a standardized major axis regression. Allometries were identified. *N*. *apicalis* and *N*. *commutata* shared a common slope between each functional brain compartment (p > 0.05, Table 2). Scaling patterns differed for each compartment; the MB (β = 0.74), AL (β = 1.36), CX (β = 1.65), and SEZ (β = 1.79) scaled allometrically (p < 0,05), while the MBP (β = 0.91), MBC (β = 0.73), MB lip (β = 0.75), MB collar (β = 0.57) and the OL (β = 1.37) scaled isometrically (p > 0.05). A significant common shift was observed for all brain compartments (p < 0.05; except for the MB collar and the OL), indicating not overlapping size differences, and despite sharing a common slope, all brain compartments (except for the MB peduncle) exhibited significant grade shifts (Table 2, Figure 2). The MB was significantly larger in *N*. *commutata* (GSI = 1.39, *p* < 0.001), a 39% larger than *N*. *apicalis*. The MBP did not differ between *N*. *apicalis* and *N*. *commutata* (GSI = 1.011, p > 0.05). The MB calyces (GSI = 1.58, *p* < 0.001) and MB lip (GSI = 1.63, *p* < 0.001) were 58% and 63% larger in *N. commutata* respectively (Table 2). The MB collar was 49% significantly larger in *N*. *apicalis* (GSI = 0.51, *p* <0.001, Figure 2, Table 2). The OL was 83% larger in *N*. *apicalis* (GSI = 0.17, *p* < 0.001; Table 2). Similarly, the AL (GSI = 0.68, *p* = 0.02), like the CX (GSI = 0.56, *p* <0.001) and the SEZ (GSI = 0.49, *p* = 0.012), were 32%, 44%, and 51% larger in *N. apicalis* (Figure 2).

**Figure 2.**
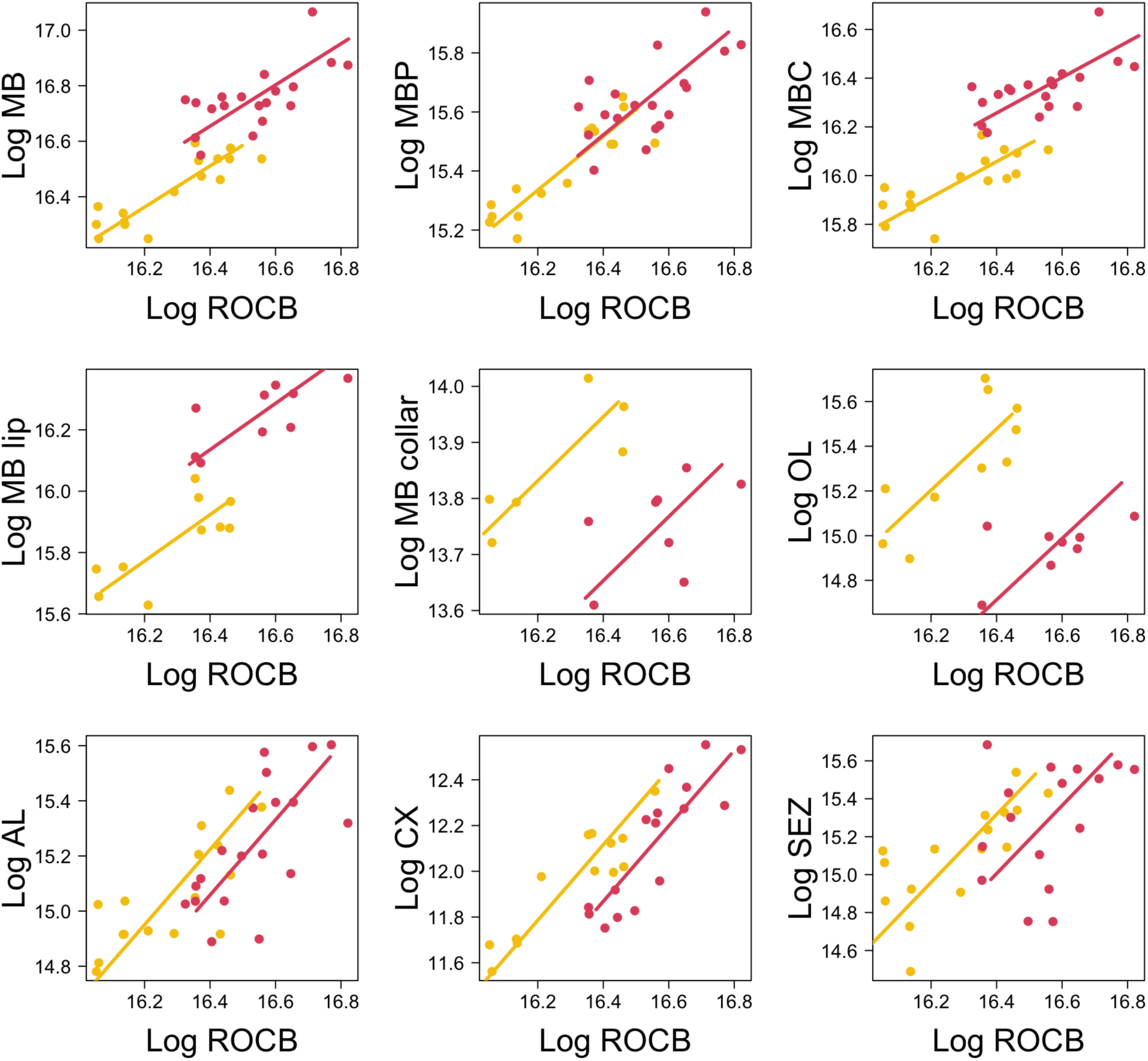
Scaling of functional brain regions relative to the remainder of the central brain volume (ROCB) in *N*. *apicalis* (yellow) and *N*. *commutata* (red) workers. Bivariate standardized major axis (SMA) regressions between the volume of log-transformed brain compartments and the ROCB.

**Table 2:** Standardized major axis regressions to test for allometry in *N*. *apicalis* and *N*. *commutata*. The volume of each functional brain compartment was compared with the ROCB volume. A grade shift index (GSI = 10^elevation*N*.^ *^commutata^*^-elevation*N*.^ *^apicalis^*) was used to compare the differences in elevation between our focal species. If GSI > 1, the brain compartment was larger in *N*. *commutata*, and if GSI < 1, the brain compartment was larger in *N*. *apicalis*. A slope index (SI = β*_N_*_. *commutata*_*/*β*_N_*_. *apicalis*_) was used to determine if both species have similar slope. If SI deviates from β = 1 then both species have different slopes (β). A log likelihood was used to test for common slope between *N*. *apicalis* and *N*. *commutata*, if p-value > 0.05 then both species share a common slope in the relationship between each functional brain compartment and the ROCB.

|  | Common slope |  | Isometry |  |  | Common shift |  | Common elevation |  |  |
| --- | --- | --- | --- | --- | --- | --- | --- | --- | --- | --- |
| Structure | Log likelihood | *P* | Log likelihood | *p* | SI | Wald test | *p* | Wald test | *p* | GSI |
| MB | 0.119 | 0.729 | 6.652 | 0.036 | 1.082 | 41.95 | <0.001 | 17.98 | <0.001 | 1.390 |
| MB peduncle | 0.01 | 0.919 | 0.778 | 0.678 | 1.023 | 22.59 | <0.001 | 0.013 | 0.909 | 1.011 |
| MB calyces | 0.046 | 0.829 | 5.401 | 0.067 | 1.058 | 53.29 | <0.001 | 25.92 | <0.001 | 1.582 |
| MB lip | 0.742 | 0.389 | 2.848 | 0.241 | 0.730 | 27.17 | <0.001 | 14.78 | <0.001 | 1.628 |
| MB collar | 0.003 | 0.955 | 4.667 | 0.097 | 0.973 | 0.345 | 0.555 | 23.34 | <0.001 | 0.509 |
| OL | 2.947 | 0.086 | 4.798 | 0.091 | 0.474 | 0.001 | 0.976 | 44.75 | <0.001 | 0.172 |
| AL | 1.189 | 0.276 | 6.658 | 0.036 | 1.320 | 13.06 | <0.001 | 5.384 | 0.020 | 0.683 |
| CX | 3.422 | 0.064 | 21.96 | <0.001 | 1.419 | 8.322 | 0.0039 | 15.75 | <0.001 | 0.565 |
| SEZ | 0.801 | 0.371 | 13.98 | <0.001 | 1.328 | 11.57 | <0.001 | 6.305 | 0.012 | 0.487 |

### Compound eye structure and optic lobe size and structure

To examine differences in eye structure and OL investment and their socioecological correlates, we analyzed the number and size of ommatidia, as well as the volume of the OL and its subregions. We found significant differences in eye structure (Figure 3): Ommatidia diameter and eye surface area were significantly larger in *N*. *commutata* (*p* < 0.001) but ommatidia number/eye surface area was significantly greater in *N*. *apicalis* (*p* < 0.001). We found no significant difference in the number of ommatidia (*p* = 0.12). The relative size of the OL (to ROCB volume) was larger in *N*. *apicalis* workers (*p* < 0.001), and a larger lamina (*p* = 0.0013) was found in *N*. *apicalis* workers; OL medulla size was not significantly different (*p* = 0.97) and the lobula was significantly larger (*p* = 0.0068) in *N*. *commutata* workers (Figure 4).

**Figure 3.**
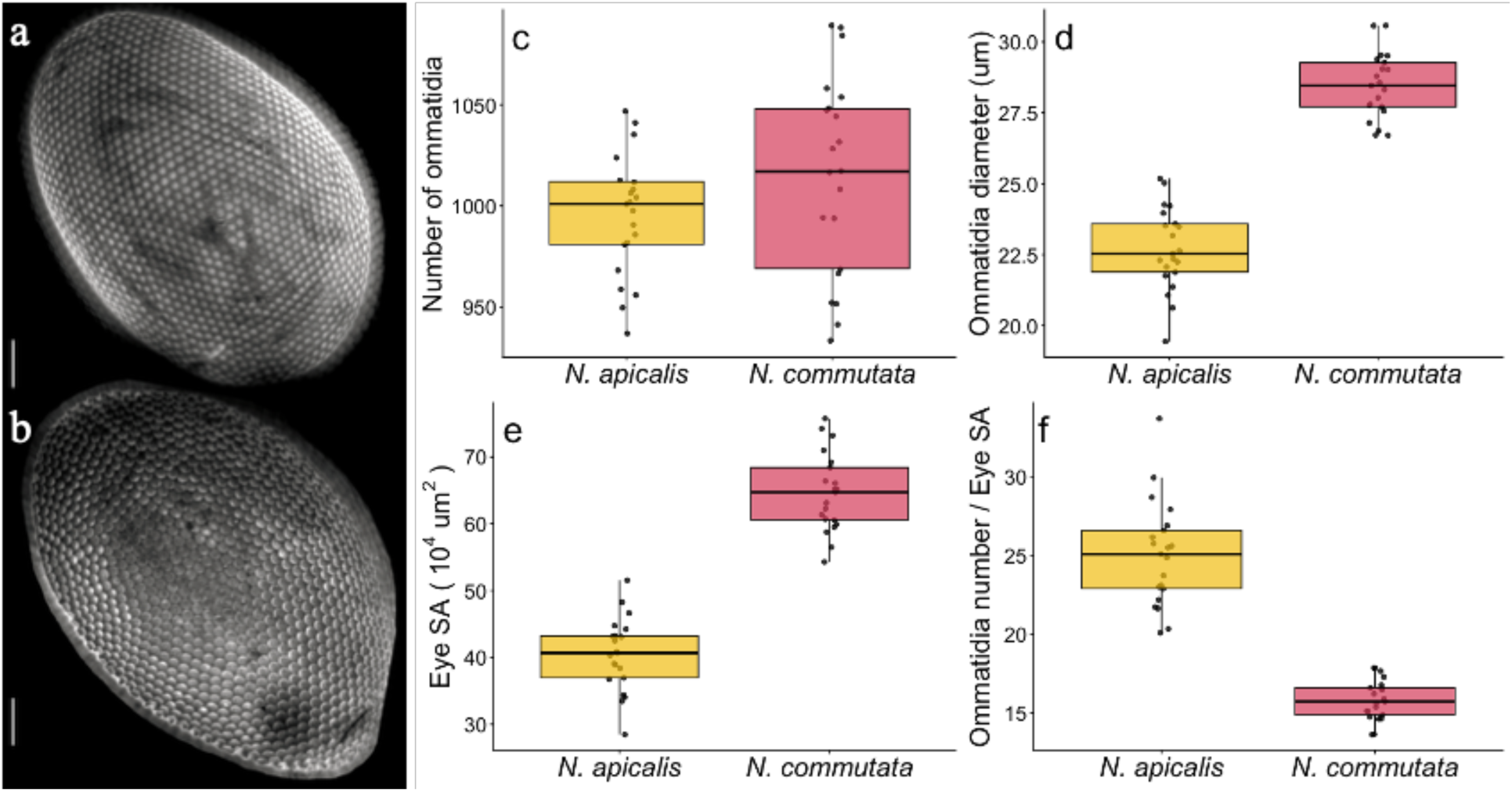
Interspecific comparison of the eye structure in two *Neoponera* ant species. Left panel: Confocal images of the eye of *N*. *apicalis* (a) and *N*. *commutata* (b). Scale bars = 100μm. Right panel: Number of ommatidia (c), ommatitida diameter (d), eye surface area [SA] (e), ommatidia density (f).

**Figure 4.**
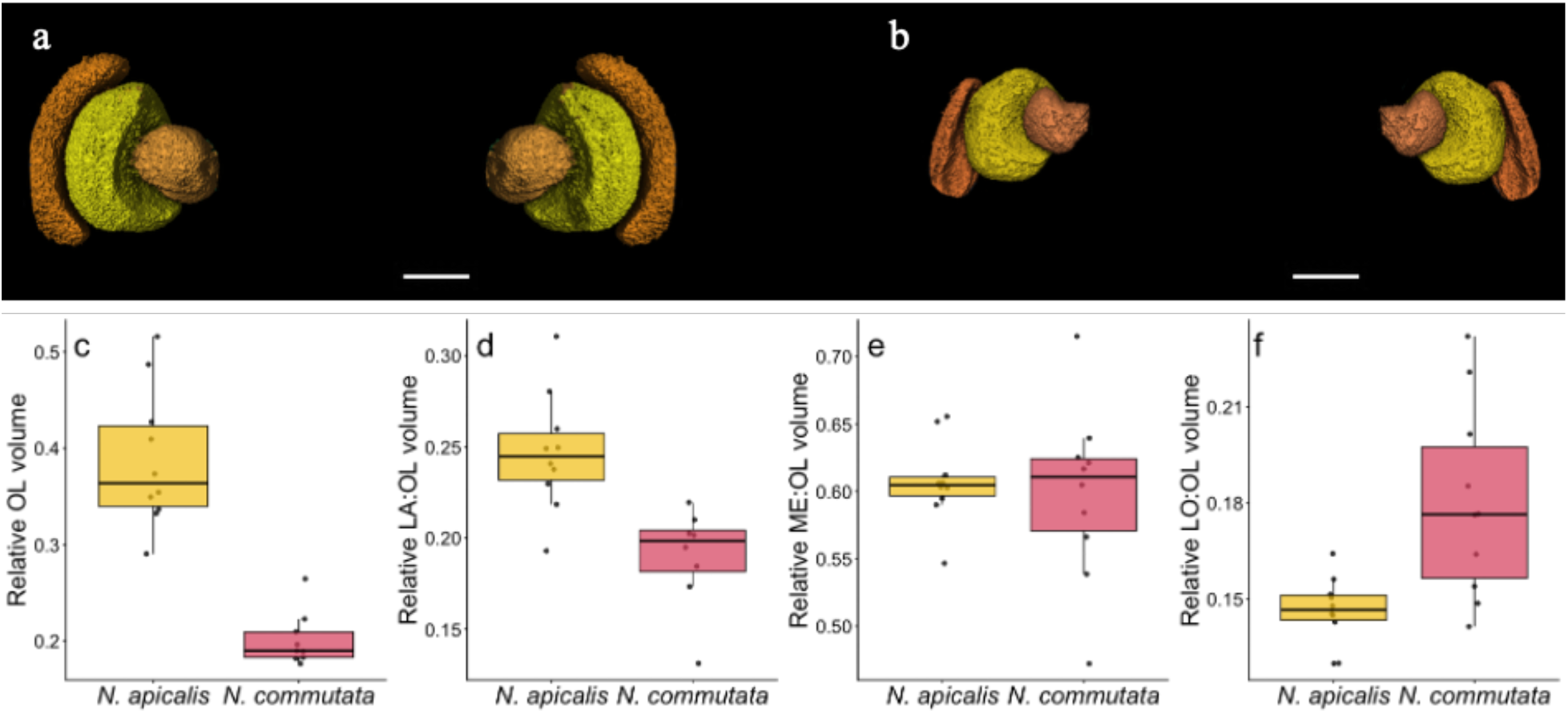
Interspecific comparison of the relative eye investment in two *Neoponera* ant species. Upper panel: 3D-reconstructions of the OL and its subcompartments in *N*. *apicalis* (a) and *N*. *commutata* (b). Scale bars = 50μm. Lower panel: OL relative investment (c) and relative investment of each OL subcompartment (relative to the OL volume), lamina [LA] (d), medulla [ME] (e) and lobula [LO] (f).

### Antennal lobe relative size; glomeruli number and size

To identify differences in olfactory processing, we compared the relative investment of the AL, and the number and size of AL glomeruli. We found no significant differences in relative investment between *N*. *apicalis* and *N*. *commutata* (*p* = 0.46) but *N. apicalis* had a significant larger number of glomeruli (*p* = 0.002), whereas the glomeruli mean volume was larger (*p* < 0.001) in *N*. *commutata* workers (Figure 5).

**Figure 5.**
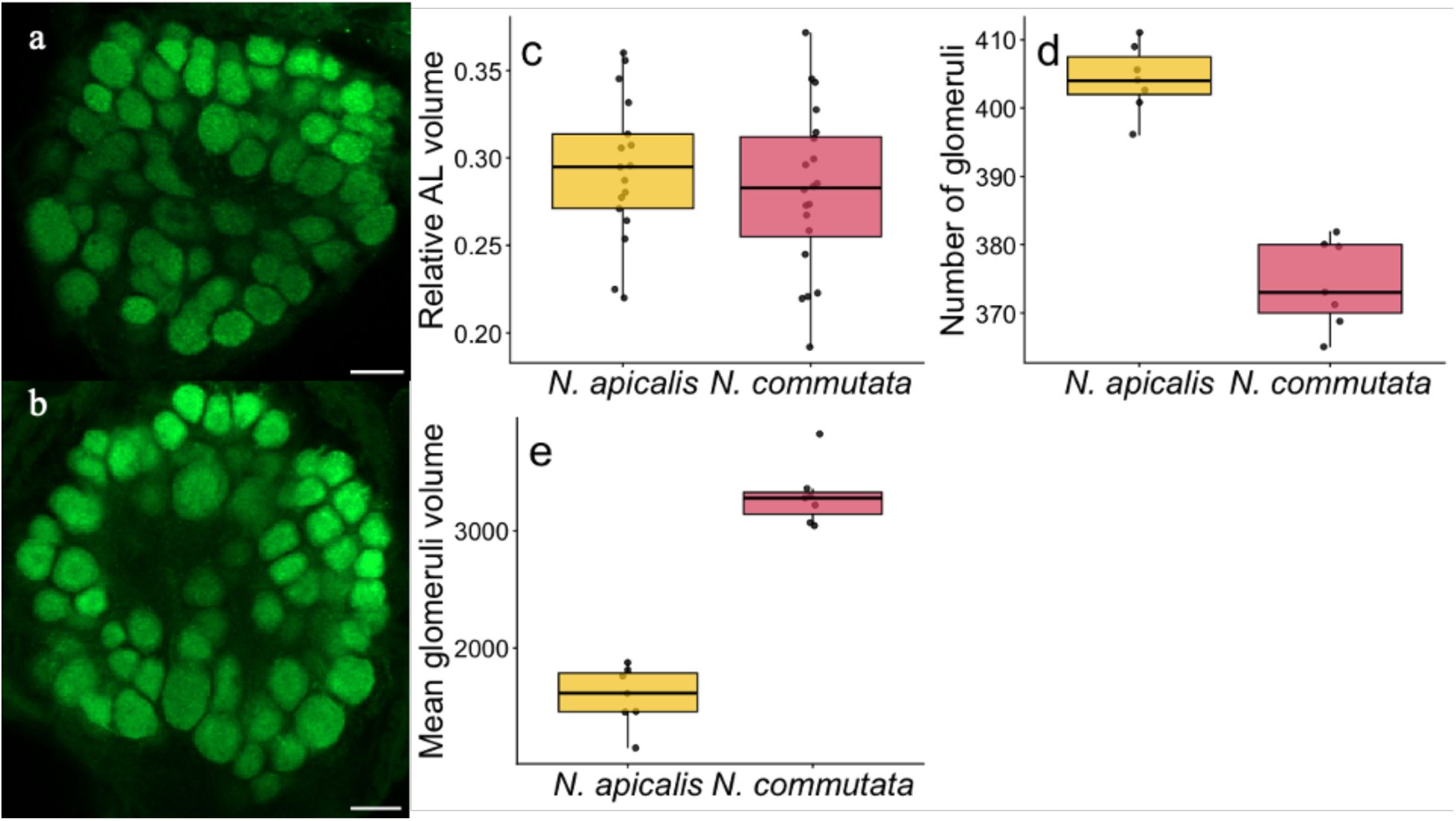
Interspecific comparison of the relative AL investment, glomeruli size and number in two *Neoponera* ant species. Left panel: Confocal images of the AL of *N*. *apicalis* (a) and *N*. *commutata* (b). Scale bars = 50μm. Left panel: AL relative size (a), AL glomeruli number (b) and size (c).

### Mushroom body relative size and structure

To determine if changes in diet and social complexity influenced higher-order processing, we compared the relative size of the MB and its subregions. The MB calyces (*p* = 0.0019) and the MB lip (*p* = 0.0101) were significantly larger in *N*. *commutata*, whereas the MB collar was significantly larger (*p* = 0.0006) in *N*. *apicalis* workers (Figure 6). The MBP was not significantly different between species (p > 0.05).

**Figure 6.**
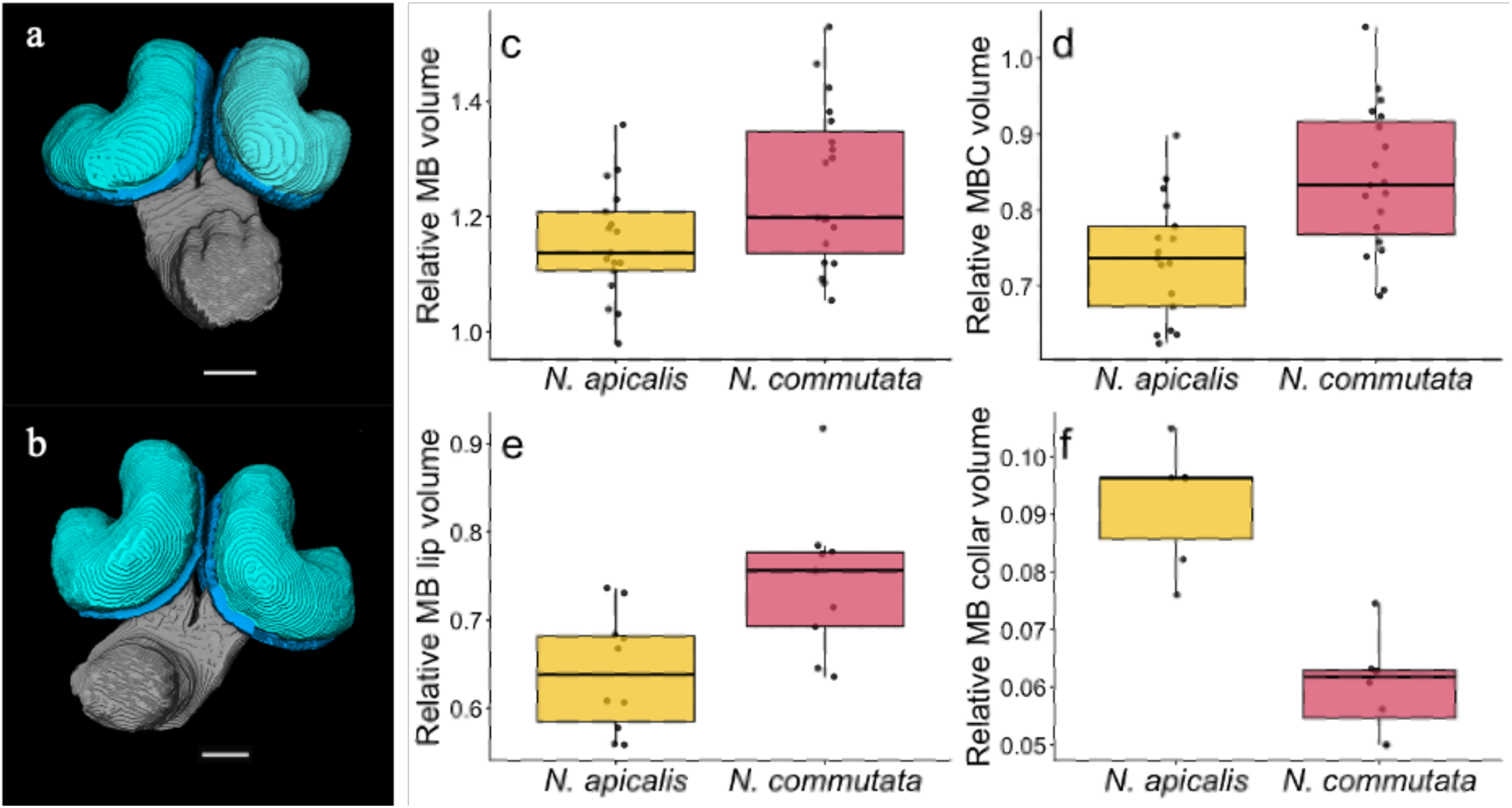
Interspecific comparison of the relative MB investment and its subcompartments in two *Neoponera* ant species. Right panel: 3-D reconstructions of the MB of *N*. *apicalis* (a) and *N*. *commutata* (b). Scale bars = 50μm. Left panel: Relative size of the mushroom body (c), MB calyces (d), MB lip (e) and MB collar (f).

### Microglomeruli density and size

To determine if variation in navigational and chemical communication demands were associated with MG structure, we measured MG density and size in the MB and found a significant higher density of MG in the MB lip (*p* < 0.001) and MB collar (*p* = 0.0056) in *N*. *apicalis*. However, the MG was significantly larger in *N*. *commutata* in both the MB lip (*p* = 0.034) and MB collar (*p* = 0.0372; Figure 7).

**Figure 7.**
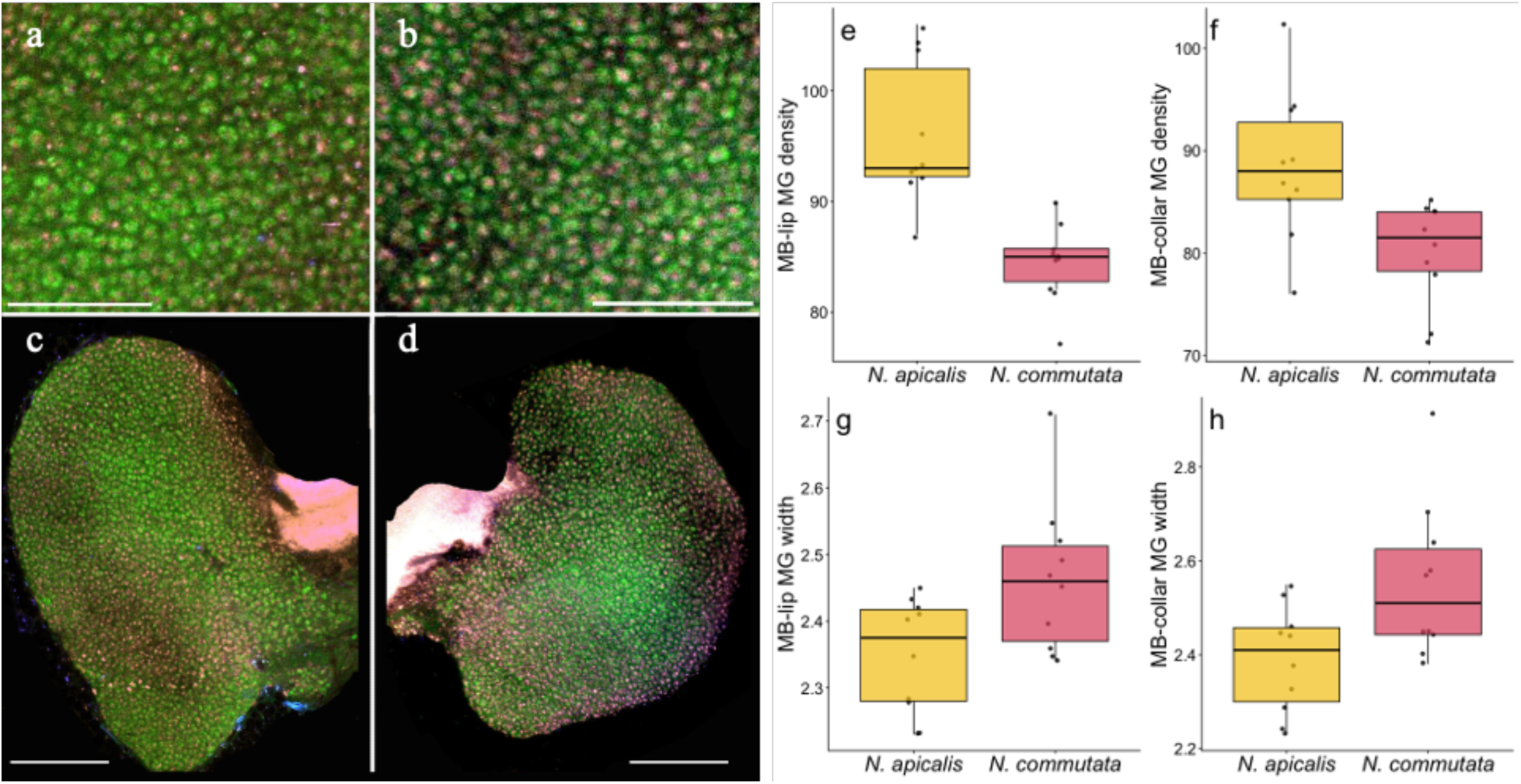
Interspecific comparison of the MB microglomeruli (MG) in two *Neoponera* ant species. Left panel: Confocal images of the MG and the MB calix of *N*. *apicalis* (a, c) and *N*. *commutata* (b, d) workers. Scale bars = 25μm (a,b), and 50μm (c,d). Right panel: MG number (e, f) and size (g, h) in the MB lip and the MB collar.

### Relative size of the central complex and subesophageal ganglion

Comparison of the relative size of the CX and SEZ found the CX was significantly larger in *N*. *apicalis* (p < 0.05), whereas the SEZ was not significantly different between species.

## Discussion

We examined the evolution of eye structure, brain size and mosaicism, and synaptic organization in sociobiologically and socioecologically divergent sister species of predatory ants by evaluating brain evolution at two neurobiological levels. Brain mosaicism correlated with diet and reliance on chemical communication for foraging. Chemically coordinated foraging in *N. commutata* is more socially complex than solitary foraging. Neuropil size-scaling patterns were associated with sensory and integrative-processing demands for prey recognition, search, homing orientation and navigation, and the presence or absence of recruitment communication during foraging. We caution that detailed information on interspecific variation in home range size and worker navigational mechanisms for our study species is very limited and it is uncertain if *N*. *commutata* workers vary in scouting, raiding, defending, and within-nest activity specializations, as suggested by Schmidt and Overal (2009). The results of our interspecific comparison nevertheless indicate a role of diet and the social organization of foraging as dual selective factors in *N*. *apicalis* and *N*. *commutata* sensory system and neuroarchitectural evolution. We discuss the sensory and neuroanatomical evidence in support of this inference below.

### Conserved relative brain volume despite differences in body size

*N. apicalis* and *N*. *commutata* workers showed no significant differences in relative brain volume, indicating that despite differences in ecology, social organization and in body size (∼ 12 mm in *N*. *apicalis* vs ∼ 17 mm in *N*. *commutata* workers) the relative brain investment was conserved. Absolute brain size was larger in *N*. *commutata* than *N*. *apicalis* workers, reflecting typical brain size/body size scaling without indicating any imprint of socioecological differentiation.

### Daily activity patterns, foraging ecology, compound eyes, and optic lobe structure

The density of ommatidia, which was significantly greater in *N*. *apicalis*, correlated with larger OL lamina size, indicating higher visual resolution and acuity (Pichaud and Casares 2022). Similar ommatidial structure is characteristic of epigeic omnivorous ants (Jelley and Barden 2021) that are mainly predatory (e.g., *Gigantiops*, *Harpegnathos*, *Myrmecia*) and rely on vision (Narendra et al. 2017). Larger OL lamina also occur in *Myrmecia* (Sheehan et al 2018) and other diurnal insects (Stockl et al. 2015). In contrast, *N*. *commutata* workers had a significantly larger ommatidia diameter and eye surface area, indicating greater light sensitivity (Land 1997), consistent with diurnal and nocturnal activity and characteristic of workers foraging in variable ambient light environments (Arganda et al. 2020; Narendra et al. 2011; Yilmaz et al. 2014). The larger OL lobula size in *N*. *commutata* is found in some nocturnal insects (Immonen et al. 2017), although diurnal and nocturnal *Myrmecia* do not show significant differences in this trait (Sheehan et al. 2018). The relative size of the OL was larger in the solitary, diet generalist, and diurnal *N*. *apicalis* workers, which search for and retrieve food, apparently relying on vision Fresneau 1985), as other visual predatory ants (Gronenberg and Liebig 1999). The OL scales allometrically in leafcutter ant polymorphic workers according to the light environment of specialized task performance (Arganda et al. 2020). *N*. *apicalis* prey -small flies, beetles, and lepidopteran larvae – are unpredictably distributed in time and space and workers may search over large areas. *N*. *apicalis* workers may also feed opportunistically on termites, although they do not recruit to termite colonies (Fresneau and Dupuy 1988). In contrast, *N*. *commutata* relies on trail pheromone communication to organize cooperative foraging on *Syntermes* s*pp*. colonies. Scouts of this species are also likely to use vision when searching for termite nests and homing to organize recruitment. Both species have evolved visual systems for navigation, which like other ants, involve celestial cues, landmark panoramas, canopy orientation and path integration (Holldobler 1980; Narendra et al. 2017).

### Diet, foraging strategy, and antennal lobe structure

*N*. *apicalis* workers had significantly more AL glomeruli than *N*. *commutata* and the latter species had significantly larger average glomeruli volume. Glomeruli number and size are associated with prey detection and discrimination (Laissue and Vosshall 2008), which in *Neoponera* are adaptively linked to dietary variation. A larger number of glomeruli may enhance odor discrimination whereas larger glomeruli could improve sensitivity to specific odors. Our findings contrast with specialist and generalist beetles that do not show significant differences in glomeruli number (Farris and Roberts 2005). A lower number of glomeruli has been found in diet-specialist bees that show high olfactory sensitivity (Burger et al. 2013; Polidori et al. 2020; Ramirez et al. 2023). Interspecific variation in glomeruli size is mainly determined by the number of synaptic olfactory receptor neuron axons (Brown et al. 2004; Kelber et al. 2009; Lin et al. 2018). Glomeruli size is larger in honeybee foragers compared to nurses or newly emerged workers (Brown et al. 2004; Winnington et al. 1996). This increase in glomeruli volume is likely associated with structural reorganization in forager brains to transition from processing social cues and signals in nurses to identifying and processing odors of floral resources (Withers et al 1993). This appears to contrast with *N*. *commutata*, a hyperspecialist, if structural changes in the glomeruli volume correlate with greater sensitivity to specific odors.

The complexity of olfactory systems results from the interplay of the number and size of AL glomeruli that correlate with specialist or generalist diets in predatory ants. Variation in size and number of glomeruli are also present in fungus-growing ants and are associated with foraging, recruitment, and olfactory discrimination (Kelber et al. 2009; Riveros et al. 2012). In contrast, *N*. *apicalis* and *N*. *commutata* showed no significant differences in AL relative size, indicating similar scaling patterns but structural reorganization may meet the demands of processing specific olfactory cues. A larger investment in the AL is correlated with task specialization in leafcutter ants (Muratore et al. 2022) and complex social behavior in wasps (Rozanski et al. 2022). However, this does not consistently correlate with social complexity or colony size (Godfrey and Gronenberg 2021; Godfrey et al. 2021; Marty et al. 2025; Ott and Rogers 2010; Riveros et al. 2012). The allometrically larger AL in *N*. *apicalis* workers could be associated with their generalist diet and detection of diverse food resources. Our results therefore show that AL organization in *Neoponera* is driven by changes in diet that influence solitary or cooperative foraging behavior and social signalling.

### Diet, group raiding behavior, mushroom body size, and synaptic structure

Workers of the hyperspecialist *N*. *commutata* had significantly larger MB than *N. apicalis*, perhaps due to their pheromonally organized group predation and nomadism. As in other ponerines of the *laevigata* group (Holldobler et al. 1996; Holldobler and Traniello 1980), *N*. *commutata* scouts search for termite colonies, home, and lay a pheromone trail to recruit nestmates. They also relocate their nest, seemingly with pheromonal control, although additional cues such as the position of the sun, skylight polarization, geomagnetic field, and terrestrial landmarks may be important as well (Acosta-Avalos et al. 2001). The size of the MB lip and collar varies with olfactory and visual demands; greater dependence on olfaction correlates with larger MB lip size, whereas the MB collar is larger in vision-dependent species (Ehmer and Gronenberg 2004; Gronenberg and Holldobler 1999). *N commutata* workers have a larger MB lip whereas *N*. *apicalis* workers have a larger collar in association with their olfactory and visual processing requirements, respectively. These findings suggest that the relative sizes of olfactory and visual input regions are influenced by diet and chemically organized group raids, as well as activity patterns (Bouchebti and Arganda 2020).

*N*. *apicalis* workers had significantly greater MG density in both the MB lip and collar, whereas *N*. *commutata* had significantly larger MG diameter. Changes in the MG density in the lip and collar have been associated with visual and olfactory learning (Gronenberg 2001; Habenstein et al. 2020; Hourcade et al. 2010). MG size is mechanistically correlated with pheromone responsiveness (Falibene et al. 2015; Kamhi et al. 2017). Larger MG appear to correlate with an increase in synapse number/axonal bouton and vesicle number that could be associated with enhanced responsiveness to chemical signals and cues (Kamhi et al 2017; Seid et al 2005). Workers of *N*. *apicalis* exhibit sector fidelity within a radius of 20-30 meters around the nest, and increase their foraging distance with experience or age (Fresneau 1985). Desert ants (*Cataglyphis*) also show sector fidelity and navigate primarily using visual cues, path integration, and landmarks (Buchkremer and Reinhold 2008; Wehner 2003). These behavioral processes are underpinned by MB synaptic organization (Grob et al. 2019; Rossler 2019; Stieb et al. 2011). *Cataglyphis* rely more on vision than on olfaction and have higher MG density in both the lip and the collar (Groh and Rossler 2011), as in *N*. *apicalis*. A higher density of MG is correlated with diet in generalist scarab beetles (Farris and Roberts 2005). Our MG results contrast with those of two ant species that differ in social complexity: workers of the weaver ant *Oecophylla smaragdina*, a socially complex species, show greater MG density in both the MB lip and collar, whereas workers of the socially basic *Formica subsericea* have larger MG diameter. Additionally, major workers of *O*. *smaragdina* have higher MG density in the MB lip, which is associated with their large pheromonal communication repertoire, and larger MG in older workers, which may relate to an increased olfactory responsiveness (Kamhi et al. 2017). Larger MG size in the *N*. *commutata* MB lip could be explained by their chemically organized raiding behavior; larger MG in the MB collar may correlate with their diurnal and nocturnal foraging activity and the need to integrate directional information from celestial and canopy cues.

### Diet and the size of the central complex and subesophageal zone

We predicted that *N*. *apicalis* would have a larger CX due to their purported visually based navigation, in contrast to the use of olfaction in the pheromonally organized foraging of *N*. *commutata. N*. *apicalis* has an allometrically larger CX, suggesting a greater need for visual memory (e.g., Collet et al. 2025; Rossler 2019). Similarly, workers of *N*. *apicalis* had a larger relative CX size, perhaps due to navigational requirement. Our results show that the SEZ is allometrically larger in *N*. *apicalis*. This could be associated with their generalist diet, and the need to process more diverse gustatory information and more precise and efficient motor control of the mouthparts (Wright 2016; Schwarz et al. 2017). However, relative SEZ investment did not change among both species, indicating similar demands during foraging.

### Conclusions and broader implications for brain evolution in carnivorous species

Our comparative study of *N*. *apicalis* and *N*. *commutata* indicate social organization during foraging and dietary variation have selected for sensory systems and macroscopic and cellular brain structure that represent adaptations to prey distribution patterns and foraging strategy. The individual task components and behavioral capabilities required for solitary hunting and prey capture appear to demand visual and olfactory acuity to recognize prey types and perhaps learn to increase efficiency to subdue prey by the kinesthetic coordination of the gaster, sting, legs, and mandibular movements. Selection for these abilities may be further enhanced in *N. apicalis* due to its small colony size and thus the nutritional benefits and colony-level need for individual foraging success. Behaviors required for foraging organization in *N. commutata* may overlap in aspects of worker navigation during food search, subduing prey, and homing, but such selection may be relaxed by the dependence on trail communication.

The inferences drawn from our study broadly inform our understanding of brain evolution in predators. Predatory arthropods like mantids show adaptive neuroanatomy (Gonzalez-Bellido et al. 2022) such as significant investment in visual systems for prey detection and capture (Althaus et al. 2024). In contrast, arachnids exhibit diverse hunting strategies, including web-building, sit-and-wait predation, and active pursuit, and rely on a combination of visual, olfactory and mechanosensory inputs (Ortega-Escobar et al. 2023). Solitary spiders may stalk their prey or construct webs and use visual and vibratory cues, respectively. Active hunting and web-building species that rely on vision have larger visual neuropils and MB but insignificant differences in the arcuate body, a brain compartment associated with mechanosensory integration (Loesel et al. 2011; Steinhoff et al. 2023). Although, females of subsocial spider species have a larger arcuate body due to higher cognitive demands of solitary foraging and the need to individually perform more tasks than social species (Caponera et al. 2021). The arcuate body appears to be less developed in solifuges, occupying 2% of the total brain volume compared to 3.1% in thelyphonids and 4.3% in spiders. Variation in this brain region is thus related to hunting strategy, prey type, or sensory ecology (Hebets et al. 2023). An increase in brain size in social canids appears due to the demands of group hunting, whereas solitary species have larger brain size due to requirements for learning and memory (Finarelli and Flynn 2009), facilitating carnivore adaptation to new environments (Michaud et al. 2022). However, the link between sociality and brain size in carnivores is unclear. Ecology seems to be the main driver of brain evolution, with dietary breath showing a negative relationship with brain size (Chambers et al 2021). Carnivores also show auditory, cerebellar and visual mosaicism; these regions scale with social behavior, ecology, home range size and daily activity patterns (Boch et al. 2024; Chambers et al. 2021; Nelson et al. 2024). Our results on predatory ant neuroarchitecture illustrate broad convergence with solitary and social carnivores in which diet, social organization, and activity pattern influence brain evolution.

## Acknowledgements

We thank Dr. Zach Coto, Dr. Mario Muscedere, Jordan Smith and Isabella Arabia for comments on the manuscript. Ant colonies were collected and maintained in the laboratory in compliance with USDA PPQ 526P-21-03079. This research was supported by NSF Grant IOS1953393 to JFAT and FONDECYT (372-2019) to FA and JFAT.

## References

Acosta-Avalos, D., D. M. Esquivel, E. Wajnberg, H. G. Lins de Barros, P. S. Oliveira, and I. Leal. 2001. “Seasonal patterns in the orientation system of the migratory ant *Pachycondyla marginata*.” Naturwissenschaften 88: 343–346.

Althaus, V., G. Exner, J. von Hadeln, U. Homberg, and R. Rosner. 2024. “Anatomical organization of the cerebrum of the praying mantis *Hierodula membranacea*.” Journal of Comparative Neurology 532, no. 3: e25607.

Amador-Vargas, S., W. Gronenberg, W. T. Wcislo, and U. Mueller. 2015. “Specialization and group size: brain and behavioural correlates of colony size in ants lacking morphological castes.” Proceedings of the Royal Society B: Biological Sciences 282, no. 1801: 20142502.

Anderson, C., and D. W. McShea. 2001. “Individual versus social complexity, with particular reference to ant colonies.” Biological reviews 76, no. 2: 211–237.

Anton, S., and U. Homberg. 1999. Antennal lobe structure. In: Hansson, B.S. (eds). “Insect olfaction”, 97–124. (Springer, Berlin, Heidelberg, 1999).

Arganda, S., A. P. Hoadley, E. S. Razdan, I. B. Muratore, and J. F. Traniello. 2020. “The neuroplasticity of division of labor: worker polymorphism, compound eye structure and brain organization in the leafcutter ant *Atta cephalotes*.” Journal of Comparative Physiology A 206: 651–662.

Azorsa, F., M. L. Muscedere, and J. F. Traniello. 2022. “Socioecology and evolutionary neurobiology of predatory ants.” Frontiers in Ecology and Evolution 9: 804200.

Barton, R. A., and P. H. Harvey. 2000. “Mosaic evolution of brain structure in mammals.” Nature 405, no. 6790: 1055–1058.

Bell-Roberts, L., J. F. Turner, G. D. Werner, P. A. Downing, L. Ross, and S. A. West. 2024. “Larger colony sizes favoured the evolution of more worker castes in ants.” Nature Ecology & Evolution 1–13.

Bertrand, O. C., Shelley, S. L., Williamson, T. E., Wible, J. R., Chester, S. G., Flynn, J. J., … and Brusatte, S. L. 2022. “Brawn before brains in placental mammals after the end-Cretaceous extinction.” Science 376, no. 6588: 80–85.

Boch, M., K. Karadachka, K. K. Loh, R. A. Benn, L. Roumazeilles, M. F. Bertelsen, … and R. B. Mars. 2024. “Comparative neuroimaging of the carnivoran brain: Neocortical sulcal anatomy.” eLife 13:RP100851.

Bouchebti, S., and S. Arganda. 2020. “Insect lifestyle and evolution of brain morphology.” Current opinion in insect science 42: 90–96.

Brown, S. M., R. M. Napper, and A. R. Mercer. 2004. “Foraging experience, glomerulus volume, and synapse number: a stereological study of the honey bee antennal lobe.” Journal of Neurobiology 60, no. 1: 40–50.

Buehlmann, C., S. Dell-Cronin, A. Diyalagoda Pathirannahelage, R. Goulard, B. Webb, J. E. Niven, and P. Graham. 2023. “Impact of central complex lesions on innate and learnt visual navigation in ants.” Journal of Comparative Physiology A 209, no. 4: 737–746.

Buchkremer, E. M., and K. Reinhold. 2008. “Sector fidelity—an advantageous foraging behavior resulting from a heuristic search strategy.” Behavioral Ecology 19, no. 5: 984–989.

Caponera, V., L. Avilés, M. Barrett, and S. O’Donnell. 2021. “Behavioral attributes of social groups determine the strength and direction of selection on neural investment.” Frontiers in Ecology and Evolution 9: 733228.

Chambers, H. R., S. A. Heldstab, and S. J. O’Hara. 2021. “Why big brains? A comparison of models for both primate and carnivore brain size evolution.” PLoS One 16, no: 12: e0261185.

Chittka, L., and J. Niven. 2009. “Are bigger brains better?.” Current biology 19, no. 21: R995–R1008.

Clutton-Brock, T. H., and P. H. Harvey. 1980. “Primates, brains and ecology.” Journal of zoology 190, no. 3: 309–323.

Collett, T., P. Graham, and S. Heinze. 2025. “The neuroethology of ant navigation.” Current Biology 35, no. 3: R110–R124.

Couto, A., F. J. Young, D. Atzeni, S. Marty, L. Melo-Flórez, L. Hebberecht, … and S. H. Montgomery. 2023. “Rapid expansion and visual specialisation of learning and memory centres in the brains of Heliconiini butterflies.” Nature Communications 14, no. 1: 4024.

DeCasien, A. R., and J. P. Higham. 2019. “Primate mosaic brain evolution reflects selection on sensory and cognitive specialization.” Nature Ecology & Evolution 3, no. 10: 1483–1493.

DeCasien, A. R., S. A. Williams, and J. P. Higham. 2017. “Primate brain size is predicted by diet but not sociality.” Nature ecology & evolution 1, no. 5: 0112.

Dejean, A., and J. P. Lachaud. 2011. “The hunting behavior of the African ponerine ant *Pachycondyla pachyderma*.” Behavioural processes 86, no. 2: 169–173.

Downing, H. 1978. “Foraging and migratory behavior of the ponerine ant Termitopone laevigata.” BA thesis, Smith College, Northampton, Massachusetts.

Dunbar, R. I. 1998. “The social brain hypothesis.” *Evolutionary Anthropology*: Issues, News, and Reviews: Issues, News, and Reviews, 6, no. 5: 178–190.

Dunbar, R. I., and S. Shultz. 2017. “Why are there so many explanations for primate brain evolution?.” Philosophical Transactions of the Royal Society B: Biological Sciences 372, no. 1727: 20160244.

Dyer, A. G., A. C. Paulk, and D.H. Reser. 2011. “Colour processing in complex environments: insights from the visual system of bees.” Proceedings of the Royal Society B: Biological Sciences 278: 952–959.

Ehmer, B., and W. Gronenberg. 2004. Mushroom body volumes and visual interneurons in ants: comparison between sexes and castes. Journal of Comparative Neurology 469, no. 2: 198–213.

Fahrbach, S. E. 2006. “Structure of the mushroom bodies of the insect brain.” Annual review of entomology 51, no. 1: 209–232.

Falibene, A., F. Roces, and W. Rössler. 2015. “Long-term avoidance memory formation is associated with a transient increase in mushroom body synaptic complexes in leaf-cutting ants.” Frontiers in behavioral neuroscience 9: 84.

Farris, S. M. 2016. “Insect societies and the social brain.” Current opinion in insect science 15: 1–8.

Farris, S. M., and N. S. Roberts. 2005. “Coevolution of generalist feeding ecologies and gyrencephalic mushroom bodies in insects.” Proceedings of the National Academy of Sciences 102, no. 48: 17394–17399.

Farris, S. M., and S. Schulmeister. 2011. “Parasitoidism, not sociality, is associated with the evolution of elaborate mushroom bodies in the brains of hymenopteran insects.” Proceedings of the Royal Society B: Biological Sciences 278, no. 1707: 940–951.

Finarelli, J. A., and J. J. Flynn. 2009. “Brain-size evolution and sociality in Carnivora.” Proceedings of the National Academy of Sciences 106, no. 23: 9345–9349.

Fresneau, D. 1985. “Individual foraging and path fidelity in a ponerine ant.” Insectes sociaux 32, no. 2: 109–116.

Fresneau, D., and P. Dupuy. 1988. “A study of polyethism in a ponerine ant: *Neoponera apicalis* (Hymenoptera, Formicidae).” Animal Behaviour 36, no. 5: 1389–1399.

Godfrey, R. K., and W. Gronenberg. 2019. “Brain evolution in social insects: advocating for the comparative approach.” Journal of Comparative Physiology A 205: 13–32.

Godfrey, R. K., J. T. Oberski, T. Allmark, C. Givens, J. Hernandez-Rivera, and W. Gronenberg. 2021. “Olfactory system morphology suggests colony size drives trait evolution in odorous ants (Formicidae: Dolichoderinae).” Frontiers in Ecology and Evolution 9: 733023.

Goldman-Huertas, B., R. F. Mitchell, R. T. Lapoint, C. P. Faucher, J. G. Hildebrand, and N. K. Whiteman. 2015. “Evolution of herbivory in Drosophilidae linked to loss of behaviors, antennal responses, odorant receptors, and ancestral diet.” Proceedings of the National Academy of Sciences 112, no. 10: 3026–3031.

Gonzalez-Bellido, P. T., J. Talley, and E. K. Buschbeck. 2022. “Evolution of visual system specialization in predatory arthropods.” Current Opinion in Insect Science 52: 100914.

Gordon, D. G., I. Ilieş, and J. F. Traniello. 2017. “Behavior, brain, and morphology in a complex insect society: trait integration and social evolution in the exceptionally polymorphic ant *Pheidole rhea*.” Behavioral Ecology and Sociobiology 71: 1–13.

Gordon, D. G., A. Zelaya, K. Ronk, and J. F. Traniello. 2018. “Interspecific comparison of mushroom body synaptic complexes of dimorphic workers in the ant genus *Pheidole*.” Neuroscience Letters 662: 110–114.

Gordon, D. G., A. Zelaya, I. Arganda-Carreras, S. Arganda, and J. F. Traniello. 2019. “Division of labor and brain evolution in insect societies: Neurobiology of extreme specialization in the turtle ant *Cephalotes varians*.” PloS one 14, no. 3: e0213618.

Graham, P., and A. Philippides. 2017. “Vision for navigation: what can we learn from ants?.” Arthropod Structure & Development 46, no. 5: 718–722.

Grob, R., P. N. Fleischmann, and W. Rössler. 2019. “Learning to navigate–how desert ants calibrate their compass systems.” Neuroforum 25, no. 2: 109–120.

Groh, C., and W. Rössler. 2011. “Comparison of microglomerular structures in the mushroom body calyx of neopteran insects.” Arthropod structure & development 40, no. 4: 358–367.

Groh, C., Z. Lu, I. A. Meinertzhagen, and W. Rössler. 2012. “Age-related plasticity in the synaptic ultrastructure of neurons in the mushroom body calyx of the adult honeybee *Apis mellifera*.” Journal of Comparative Neurology 520: 3509–3527.

Groh, C., C. Kelber, K. Grübel, and W. Rössler. 2014. “Density of mushroom body synaptic complexes limits intraspecies brain miniaturization in highly polymorphic leaf-cutting ant workers.” Proceedings of the Royal Society B: Biological Sciences 281, no. 1785: 20140432.

Gronenberg, W., and B. Hölldobler. 1999. “Morphologic representation of visual and antennal information in the ant brain.” Journal of Comparative Neurology 412, no. 2: 229–240.

Gronenberg, W., and J. Liebig. 1999. “Smaller brains and optic lobes in reproductive workers of the ant *Harpegnathos*.” Naturwissenschaften 86: 343–345.

Gronenberg, W. 2001. “Subdivisions of hymenopteran mushroom body calyces by their afferent supply.” Journal of Comparative Neurology 435, no. 4: 474–489.

Gronenberg, W. 2008. “Structure and function of ant (Hymenoptera: Formicidae) brains: strength in numbers.” Myrmecological News 11: 25–36.

Habenstein, J., E. Amini, K. Grübel, B. El Jundi, and W. Rössler. 2020. “The brain of *Cataglyphis* ants: Neuronal organization and visual projections.” Journal of Comparative Neurology 528, no. 18: 3479–3506.

Harvey, P. H., T. H. Clutton-Brock, and G. M. Mace. 1980. “Brain size and ecology in small mammals and primates.” Proceedings of the National Academy of Sciences 77, no. 7: 4387–4389.

Hansson, B. S., and M. C. Stensmyr. 2011. “Evolution of insect olfaction.” Neuron 72, no. 5: 698–711.

Hebets, E. A., M. Oviedo-Diego, F. Cargnelutti, F. Bollatti, L. Calbacho-Rosa, C. I. Mattoni, … and A. V. Peretti. 2024. “A scientist’s guide to Solifugae: how solifuges could advance research in ecology, evolution, and behaviour.” Zoological Journal of the Linnean Society 202, no. 2: 1–27. 10.1093/zoolinnean/zlad174

Hölldobler, B. 1980. “Canopy orientation: a new kind of orientation in ants.” Science 210, no. 4465: 86–88.

Hölldobler, B., and J. F. Traniello. 1980. “The pygidial gland and chemical recruitment communication in *Pachycondyla* (= *Termitopone*) *laevigata*.” Journal of Chemical Ecology 6: 883–893.

Hölldobler, B., and E. O. Wilson. 1990. The ants. Harvard University Press.

Hölldobler, B., E. Janssen, H. J. Bestmann, F. Kern, I. R. Leal, P. S. Oliveira, and W. A. König. 1996. “Communication in the migratory termite-hunting ant *Pachycondyla* (= *Termitopone*) *marginata* (Formicidae, Ponerinae).” Journal of Comparative Physiology A 178: 47–53.

Hourcade, B., T. S. Muenz, J. C. Sandoz, W. Rössler, and J. M. Devaud. 2010. “Long-term memory leads to synaptic reorganization in the mushroom bodies: a memory trace in the insect brain?” Journal of Neuroscience 30, no. 18: 6461–6465.

Immonen, E. V., M. Dacke, S. Heinze, and B. El Jundi. 2017. “Anatomical organization of the brain of a diurnal and a nocturnal dung beetle.” Journal of Comparative Neurology 525, no. 8: 1879–1908.

Ito, K., K. Shinomiya, M. Ito, J. D. Armstrong, G. Boyan, V. Hartenstein, … and L. B. Vosshall. 2014. “A systematic nomenclature for the insect brain.” Neuron 81, no. 4: 755–765.

Iwaniuk, A. N., K. M. Dean, and J. E. Nelson. 2004. “A mosaic pattern characterizes the evolution of the avian brain.” Proceedings of the Royal Society B: Biological Sciences 271: S148–S151.

Jaffe, K., and E. Perez. 1989. “Comparative study of brain morphology in ants.” Brain, behavior and evolution 33, no. 1: 25–33.

Jelley, C., and P. Barden. 2021. “Vision-linked traits associated with antenna size and foraging ecology across ants.” Insect Systematics and Diversity 5, no. 5: 9.

Kamhi, J. F., W. Gronenberg, S. K. Robson, and J. F. Traniello. 2016. “Social complexity influences brain investment and neural operation costs in ants.” Proceedings of the Royal Society B: Biological Sciences 283, no. 1841: 20161949.

Kamhi, J. F., A. Sandridge-Gresko, C. Walker, S. K. Robson, and J. F. A. Traniello. 2017. “Worker brain development and colony organization in ants: does division of labor influence neuroplasticity?” Developmental Neurobiology 77, no. 9: 1072–1085.

Kendroud, S., A. A. Bohra, P. A. Kuert, B. Nguyen, O. Guillermin, S. G. Sprecher, … and V. Hartenstein. 2018. “Structure and development of the subesophageal zone of the Drosophila brain. II. Sensory compartments.” Journal of Comparative Neurology 526, no. 1: 33–58.

Kelber, C., W. Rössler, F. Roces, and C. J. Kleineidam. 2009. “The antennal lobes of fungus-growing ants (Attini): neuroanatomical traits and evolutionary trends.” Brain, behavior and evolution 73, no. 4: 273–284.

Laissue, P.P., and L.B. Vosshall. 2008. “The Olfactory Sensory Map in *Drosophila*. In: Technau, G.M. (eds) Brain Development in Drosophila melanogaster.” Advances in Experimental Medicine and Biology vol 628. (Springer, 2008), New York, NY.

Land, M. F. 1997. “Visual acuity in insects.” Annual review of entomology 42, no. 1: 147–177.

Lepeco, A., O. M. Meira, D. M. Matielo, C. R. Brandão, and G. P. Camacho. 2025. “A hell ant from the Lower Cretaceous of Brazil.” Current Biology 35, no. 9: 2146–2153.

Li, L., H. MaBouDi, M. Egertová, M. R. Elphick, L. Chittka, and C. J. Perry. 2017. “A possible structural correlate of learning performance on a colour discrimination task in the brain of the bumblebee.” Proceedings of the Royal Society B: Biological Sciences 284, no. 1864: 20171323.

Lihoreau, M., T. Latty, and L. Chittka. 2012. “An exploration of the social brain hypothesis in insects.” Frontiers in physiology 3: 442.

Lihoreau, M., T. Dubois, T. Gomez-Moracho, S. Kraus, C. Monchanin, and C. Pasquaretta. 2019. “Putting the ecology back into insect cognition research.” In Advances in insect physiology (Vol. 57, pp. 1-25). Academic Press.

Lin, T., C. Li, J. Liu, B. H. Smith, H. Lei, and X. Zeng. 2018. “Glomerular organization in the antennal lobe of the oriental fruit fly *Bactrocera dorsalis*.” Frontiers in Neuroanatomy 12: 71.

Loesel, R., E. A. Seyfarth, P. Bräunig, and H. J. Agricola. 2011. “Neuroarchitecture of the arcuate body in the brain of the spider *Cupiennius salei* (Araneae, Chelicerata) revealed by allatostatin-, proctolin-, and CCAP-immunocytochemistry and its evolutionary implications.” Arthropod Structure & Development 40: 210–220.

Maguire, E. A., D. G. Gadian, I. S. Johnsrude, C. D. Good, J. Ashburner, R. S. Frackowiak, and C. D. Frith. 2000. “Navigation-related structural change in the hippocampi of taxi drivers.” Proceedings of the National Academy of Sciences 97, no. 8: 4398–4403.

Marty, S., A. Couto, E. H. Dawson, N. Brard, P. d’Ettorre, S. H. Montgomery, and J. C. Sandoz. 2025. “Ancestral complexity and constrained diversification of the ant olfactory system.” Proceedings of the Royal Society B: Biological Sciences 292, no. 2045: 20250662.

Michaud, M., S. L. D. Toussaint, and E. Gilissen. 2022. “The impact of environmental factors on the evolution of brain size in carnivorans.” Communications Biology 5, no. 1: 998.

Mill, A. E. 1984. “Predation by the ponerine ant *Pachycondyla commutata* on termites of the genus *Syntermes* in Amazonian rain forest.” Journal of natural History 18, no. 3: 405–410.

Muratore, I. B., E. M. Fandozzi, and J. F. A. Traniello. 2022. “Behavioral performance and division of labor influence brain mosaicism in the leafcutter ant *Atta cephalotes*.” Journal of Comparative Physiology A 208, no. 2: 325–344.

Muscedere, M. L., and J. F. Traniello. 2012. “Division of labor in the hyperdiverse ant genus *Pheidole* is associated with distinct subcaste-and age-related patterns of worker brain organization.” PLoS One 7, no. 2: e31618.

Muscedere, M. L., W. Gronenberg, C. S. Moreau, and J. F. Traniello. 2014. “Investment in higher order central processing regions is not constrained by brain size in social insects.” Proceedings of the Royal Society B: Biological Sciences 281, no. 1784: 20140217.

Narendra, A., S. F. Reid, B. Greiner, R. A. Peters, J. M. Hemmi, W. A. Ribi, and J. Zeil. 2011. “Caste-specific visual adaptations to distinct daily activity schedules in Australian *Myrmecia* ants.” Proceedings of the Royal Society B: Biological Sciences 278, no. 1709: 1141–1149.

Narendra, A., J. F. Kamhi, and Y. Ogawa. 2017. “Moving in dim light: behavioral and visual adaptations in nocturnal ants.” Integrative and comparative biology 57, no. 5: 1104–1116.

Nelson, J., E. M. Woeste, K. Oba, K. Bitterman, B. K. Billings, J. Sacco, … and M. A. Spocter. 2024. “Neuropil variation in the prefrontal, motor, and visual cortex of six felids.” Brain Behavior and Evolution 99, no. 1: 25–44.

Nilsson, D. E., and A. Kelber. 2007. “A functional analysis of compound eye evolution.” Arthropod structure & development 36, no. 4: 373–385.

O’Donnell, S., M. R. Clifford, S. DeLeon, C. Papa, N. Zahedi, and S. J. Bulova. 2013. “Brain size and visual environment predict species differences in paper wasp sensory processing brain regions (Hymenoptera: Vespidae, Polistinae).” Brain Behavior and Evolution 82, no. 3: 177–184.

O’Donnell, S., S. J. Bulova, S. DeLeon, P. Khodak, S. Miller, and E. Sulger. 2015. “Distributed cognition and social brains: reductions in mushroom body investment accompanied the origins of sociality in wasps (Hymenoptera: Vespidae).” Proceedings of the Royal Society B: Biological Sciences 282, no. 1810: 20150791.

O’Donnell, S., S. Bulova, M. Barrett, and C. von Beeren. 2018. “Brain investment under colony-level selection: soldier specialization in Eciton army ants (Formicidae: Dorylinae).” BMC zoology 3: 1–6.

Ortega-Escobar, J., E. A. Hebets, V. P. Bingman, D. D. Wiegmann, and D. D. Gaffin. 2023. “Comparative biology of spatial navigation in three arachnid orders (Amblypygi, Araneae, and Scorpiones).” Journal of Comparative Physiology A 209, no. 4: 747–779.

Ott, S. R., and S. M. Rogers. 2010. “Gregarious desert locusts have substantially larger brains with altered proportions compared with the solitarious phase.” Proceedings of the Royal Society B: Biological Sciences 277, no. 1697: 3087–3096.

Pichaud, F., and F. Casares. 2022. “Shaping an optical dome: The size and shape of the insect compound eye.” In Seminars in cell & developmental biology (Vol. 130, pp. 37-44). Academic Press.

Perrichot, V., B. Wang, and M. S. Engel. 2016. “Extreme morphogenesis and ecological specialization among Cretaceous basal ants.” Current Biology 26, no. 11: 1468–1472.

Powell, L. E., K. Isler, and R. A. Barton. 2017. “Re-evaluating the link between brain size and behavioural ecology in primates.” Proceedings of the Royal Society B: Biological Sciences 284, no. 1865: 20171765.

Riveros, A. J., M. A. Seid, and W. T. Wcislo. 2012. “Evolution of brain size in class-based societies of fungus-growing ants (Attini).” Animal behaviour 83, no. 4: 1043–1049.

Rosati, A. G. 2017. “Foraging cognition: reviving the ecological intelligence hypothesis.” Trends in cognitive sciences 21, no. 9: 691–702.

Rosner, R., J. von Hadeln, T. Salden, and U. Homberg. 2017. “Anatomy of the lobula complex in the brain of the praying mantis compared to the lobula complexes of the locust and cockroach.” Journal of Comparative Neurology 525, no. 10: 2343–2357.

Rössler, W. 2019. “Neuroplasticity in desert ants (Hymenoptera: Formicidae)–importance for the ontogeny of navigation.” Myrmecological News 29.

Rössler, W. 2023. “Multisensory navigation and neuronal plasticity in desert ants.” Trends in Neurosciences 46, no. 6: 415–417.

Rössler, W., R. Grob, and P. N. Fleischmann. 2023. “The role of learning-walk related multisensory experience in rewiring visual circuits in the desert ant brain.” Journal of Comparative Physiology A 209, no. 4: 605–623.

Rozanski, A. N., A. Cini, T. E. Lopreto, K. M. Gandia, M. E. Hauber, R. Cervo, and F. M. Uy. 2022. “Differential investment in visual and olfactory brain regions is linked to the sensory needs of a wasp social parasite and its host.” Journal of Comparative Neurology 530, no. 4: 756–767.

Sayol, F., M. Á. Collado, J. Garcia-Porta, M. A. Seid, J. Gibbs, A. Agorreta, … and I. Bartomeus. 2020. “Feeding specialization and longer generation time are associated with relatively larger brains in bees.” Proceedings of the Royal Society B: Biological Sciences 287, no. 1935: 20200762.

Schmidt, J. O., and W. L. Overal. 2009. “Venom and task specialization in *Termitopone commutata* (Hymenoptera: Formicidae).” Journal of Hymenoptera Research 18: 361–367.

Schmidt, C. A., and S. O. Shattuck. 2014. “The higher classification of the ant subfamily Ponerinae (Hymenoptera: Formicidae), with a review of ponerine ecology and behavior.” Zootaxa 3817, no. 1: 1–242.

Schwarz, O., A. A. Bohra, X. Liu, H. Reichert, K. VijayRaghavan, and J. Pielage. 2017. “Motor control of *Drosophila* feeding behavior.” Elife 6: e19892.

Seid, M. A., K. M. Harris, and J. F. Traniello. 2005. “Age-related changes in the number and structure of synapses in the lip region of the mushroom bodies in the ant *Pheidole dentata*.” Journal of Comparative Neurology 488, no. 3: 269–277.

Sheehan, Z. B., J. F. Kamhi, M. A. Seid, and A. Narendra. 2019. “Differential investment in brain regions for a diurnal and nocturnal lifestyle in Australian *Myrmecia* ants.” Journal of Comparative Neurology 527, no. 7: 1261–1277.

Shultz, S., and R. I. M. Dunbar. 2010. “Species differences in executive function correlate with hippocampus volume and neocortex ratio across nonhuman primates.” Journal of Comparative Psychology 124, no. 3: 252.

Shultz, S., and R. I. Dunbar. 2022. “Socioecological complexity in primate groups and its cognitive correlates.” Philosophical Transactions of the Royal Society B 377, no. 1860: 20210296.

Smith, E. J., J. Vizueta, M. A. Younger, S. P. Mullen, and J. F. Traniello. 2023. “Dietary diversity, sociality, and the evolution of ant gustation.” Frontiers in Ecology and Evolution 11: 1175719.

Steinhoff, P. O., S. Harzsch, and G. Uhl. 2023. “Comparative neuroanatomy of the central nervous system in web-building and cursorial hunting spiders.” Journal of Comparative Neurology 532, no. 2: e25554.

Stieb, S. M., T. S. Muenz, R. Wehner, and W. Rössler. 2010. “Visual experience and age affect synaptic organization in the mushroom bodies of the desert ant *Cataglyphis fortis*.” Developmental Neurobiology 70, no. 6: 408–423.

Stieb, S. M., A. Hellwig, R. Wehner, and W. Rössler. 2012. “Visual experience affects both behavioral and neuronal aspects in the individual life history of the desert ant *Cataglyphis fortis*.” Developmental neurobiology 72, no. 5: 729–742.

Stöckl, A. L., W. A. Ribi, and E. J. Warrant. 2016. “Adaptations for nocturnal and diurnal vision in the hawkmoth lamina.” Journal of Comparative Neurology 524, no. 1: 160–175.

Strausfeld, N. J. 1989. “Beneath the compound eye: neuroanatomical analysis and physiological correlates in the study of insect vision.” *Facets of Vision*. Springer, Berlin, Heidelberg, pp 317–359

Strausfeld, N. J. 2012. “Arthropod brains: evolution, functional elegance, and historical significance.” *Harvard University Press*.

Stringer, C., T. Wang, M. Michaelos, and M. Pachitariu. 2021. “Cellpose: a generalist algorithm for cellular segmentation.” Nature methods 18, 1: 100–106.

Sukhum, K. V., J. Shen, and B. A. Carlson. 2018. “Extreme enlargement of the cerebellum in a clade of teleost fishes that evolved a novel active sensory system.” Current Biology 28, no. 23: 3857–3863.

Traniello, J. F., T. A. Linksvayer, and Z. N. Coto. 2022. “Social complexity and brain evolution: insights from ant neuroarchitecture and genomics.” Current opinion in insect science 53: 100962.

Valadares, L., H. G. Rödel, P. d’Ettorre, and J. C. Sandoz. 2026. “Neural investment patterns reflect task specialization in the leaf-cutting ant *Acromyrmex subterraneus*.” Iscience 29.

Warton, D. I., R. A. Duursma, D. S. Falster, and S. Taskinen. 2012. “smatr 3-an R package for estimation and inference about allometric lines.” Methods in Ecology & Evolution 3, no. 2.

Wehner, R. 2003. “Desert ant navigation: how miniature brains solve complex tasks.” Journal of Comparative Physiology A 189: 579–588.

Wehner, R., T. Fukushi, and K. Isler. 2007. “On being small: brain allometry in ants.” Brain Behavior and Evolution 69, no. 3: 220–228.

Winnington, A. P., R. M. Napper, and A. R. Mercer. 1996. “Structural plasticity of identified glomeruli in the antennal lobes of the adult worker honey bee.” Journal of Comparative Neurology 365, no. 3: 479–490.

Wright, G. A. 2016. “To feed or not to feed: circuits involved in the control of feeding in insects.” Current Opinion in Neurobiology 41: 87–91.

Yilmaz, A., V. Aksoy, Y. Camlitepe, and M. Giurfa. 2014. “Eye structure, activity rhythms, and visually-driven behavior are tuned to visual niche in ants.” Frontiers in behavioral neuroscience 8: 205.

